# PGE_2_ production at sites of tissue injury promotes an anti-inflammatory neutrophil phenotype and determines the outcome of inflammation resolution *in vivo*

**DOI:** 10.1101/205997

**Authors:** Catherine A. Loynes, Jou A. Lee, Anne L. Robertson, Michael JG. Steel, Felix Ellett, Yi Feng, Bruce D. Levy, Moira K Whyte, Stephen A. Renshaw

## Abstract

Neutrophils are the first immune cells recruited to a site of injury or infection, where they perform many functions. Having completed their role, neutrophils must be removed from the inflammatory site - either by apoptosis and efferocytosis or by reverse migration away from the wound - for restoration of normal tissue homeostasis. Disruption of these tightly controlled physiological processes of neutrophil removal can lead to a range of inflammatory diseases. We used an *in vivo* zebrafish model to understand the role of lipid mediator production in neutrophil removal. Following tailfin amputation in the absence of macrophages, neutrophillic inflammation does not resolve. This is due to loss of macrophage-dependent production of eicosanoid prostaglandin E2, which drives neutrophil removal via promotion of reverse migration. Knockdown of endogenous prostaglandin E synthase gene reveals PGE_2_ as essential for neutrophil inflammation resolution. Furthermore, PGE_2_ is able to signal through EP4 receptors to enhance Alox15 production, causing a switch towards anti-inflammatory eicosanoid signalling, specifically Lipoxin A_4_. Our data confirm regulation of neutrophil migration by PGE_2_ and LXA_4_ in an *in vivo* model of inflammation resolution. This pathway may contain therapeutic targets for driving inflammation resolution in chronic inflammatory disease.

## Introduction

Inflammation is a critical process that maintains normal tissue homeostasis following injury or infection, by removing potential pathogens and beginning the process of wound repair and healing. Neutrophils are the first immune cells recruited to a site of injury or infection, where they carry out a number of functions (*1*). They destroy foreign pathogens, control the spread of infection and thereby minimise tissue damage. Neutrophils engulf bacteria and cell debris at the site of inflammation through phagocytosis and release antimicrobial molecules by degranulation. This, along with the release of reactive oxygen species, aids destruction of pathogens (*2*) and permits neutrophil penetration into otherwise inaccessible tissues (*3*). Having completed their role, neutrophils must be removed from the inflammatory site for successful inflammation resolution to occur. If this tightly controlled physiological process is disrupted, failure of inflammation resolution can lead to inflammatory diseases such as chronic obstructive pulmonary disease (COPD).

There are several mechanisms by which neutrophils can be removed from sites of inflammation. Caspase-dependent neutrophil apoptosis and subsequent efferocytosis by macrophages has been shown to contribute to the successful resolution of inflammation in mouse (*4*), human (*5*) and zebrafish (*6*). Natural killer cells can also induce granulocyte apoptosis as a pro-resolving mechanism (*7*). Neutrophil apoptosis is tightly coupled to uptake by macrophages to prevent tissue damage (*8*). Macrophages phagocytose pathogens and apoptotic cells and contribute to inflammation resolution (*9*), tissue remodelling and regeneration (*9*, *10*); however, their influence on inflammation resolution is incompletely defined.

More recently, it has been demonstrated that neutrophils can also undergo reverse migration away from sites of inflammation, which in some circumstances might contribute to the dissipation of the inflammatory burden and hence to inflammation resolution (*11*, *12*). In the zebrafish, reverse migration is anti-inflammatory, and is suppressed by pro-inflammatory stimuli such as hypoxia (*13*). Variations on this process have also been seen in mouse models (*14*, *15*), and in human neutrophils (*16*, *17*). Understanding the molecular mechanisms that govern neutrophil removal, either via efferocytosis or reverse migration, is essential to developing new therapeutics for inflammation resolution.

Macrophages are well known to be key mediators in determining the outcome of the inflammatory response (*18*). Macrophages and neutrophils secrete a range of pro- and anti-inflammatory cytokines, dependent on the local inflammatory environment. Studies in mammalian systems (*19*, *20*) have shown that upon uptake of apoptotic cells, macrophages regulate pro- and anti-inflammatory cytokine production (*21*). When fed with apoptotic neutrophils, LPS-stimulated human monocyte-derived macrophages downregulate key pro-inflammatory cytokines and upregulate anti-inflammatory mediators TGFβ1, PGE_2_ and PAF.

Prostaglandins, such as PGE_2_, are not stored in cells, but rather produced following arachidonic acid metabolism and therefore can be produced by almost all cells in the body (*22*).PGE_2_ is the most abundant prostaglandin in humans (*23*) generated via the action of cyclooxygenase (COX) enzymes. While PGE_2_ is known to have pro-inflammatory properties under defined conditions (*24*), it has also been shown to have anti-inflammatory functions (*25*). In zebrafish cancer models, inhibition of PGE_2_ production led to a change in macrophage behaviour and enhanced pro-inflammatory activity (*26*), consistent with PGE_2_ directing macrophages towards an anti-inflammatory phenotype (*27*). In human peripheral blood neutrophils, PGE_2_ switches lipid mediator biosynthesis from predominantly pro-inflammatory leukotriene B_4_ (5-lipoxygenase (5-LO)-initiated pathway) to lipoxin A_4_, which reduces neutrophil infiltration into exudates (*28*). These studies support a role for PGE_2_ in influencing immune cell phenotype and led us to ask whether macrophages might be altering local lipid mediator signalling as a mechanism for influencing inflammation resolution *in vivo*.

Using a zebrafish model of *in vivo* inflammation resolution, we examined the potential role of the lipid mediator PGE_2_ in driving inflammation resolution. We demonstrate that macrophage uptake of apoptotic cells is necessary for successful inflammation resolution; PGE_2_ is produced and acts via EP4 receptors to drive inflammation resolution by reverse migration.

## Results

### Macrophage clearance of apoptotic cells is necessary for successful resolution of neutrophilic inflammation *in vivo*

Inflammation requires the presence and activity of both neutrophils and macrophages. To test whether neutrophil dynamics were influenced by the presence of macrophages, we ablated macrophages using tissue specific bacterial nitroreductase expression and treating with metronidazole as previously described (*29*, *30*). Injury was performed on 3 day post fertilisation (dpf) macrophage ablated larvae and neutrophil and macrophage numbers assessed at the time points indicated (Figure 1a). In control larvae, neutrophil numbers peaked between 4 and 6 hours post injury (hpi), with numbers returning to basal levels by 24 hpi, demonstrating spontaneous resolution of inflammation. Macrophages were recruited to the site of injury at a lower rate, with numbers plateauing at 12 hpi and remaining high until at least 24 hpi. In metronidazole treated larvae, macrophages were not seen at the wound throughout the inflammatory time course, confirming successful ablation. Neutrophil numbers at the wound site were significantly increased at 24 hpi in macrophage-depleted larvae compared to control larvae, suggesting failed inflammation resolution (Figure 1a,b). This was in marked contrast to whole body neutrophil counts, which were unaltered (Figure 1c). This demonstrates that macrophages are necessary at the site of tissue injury for normal inflammation resolution to occur.

**Figure 1.**
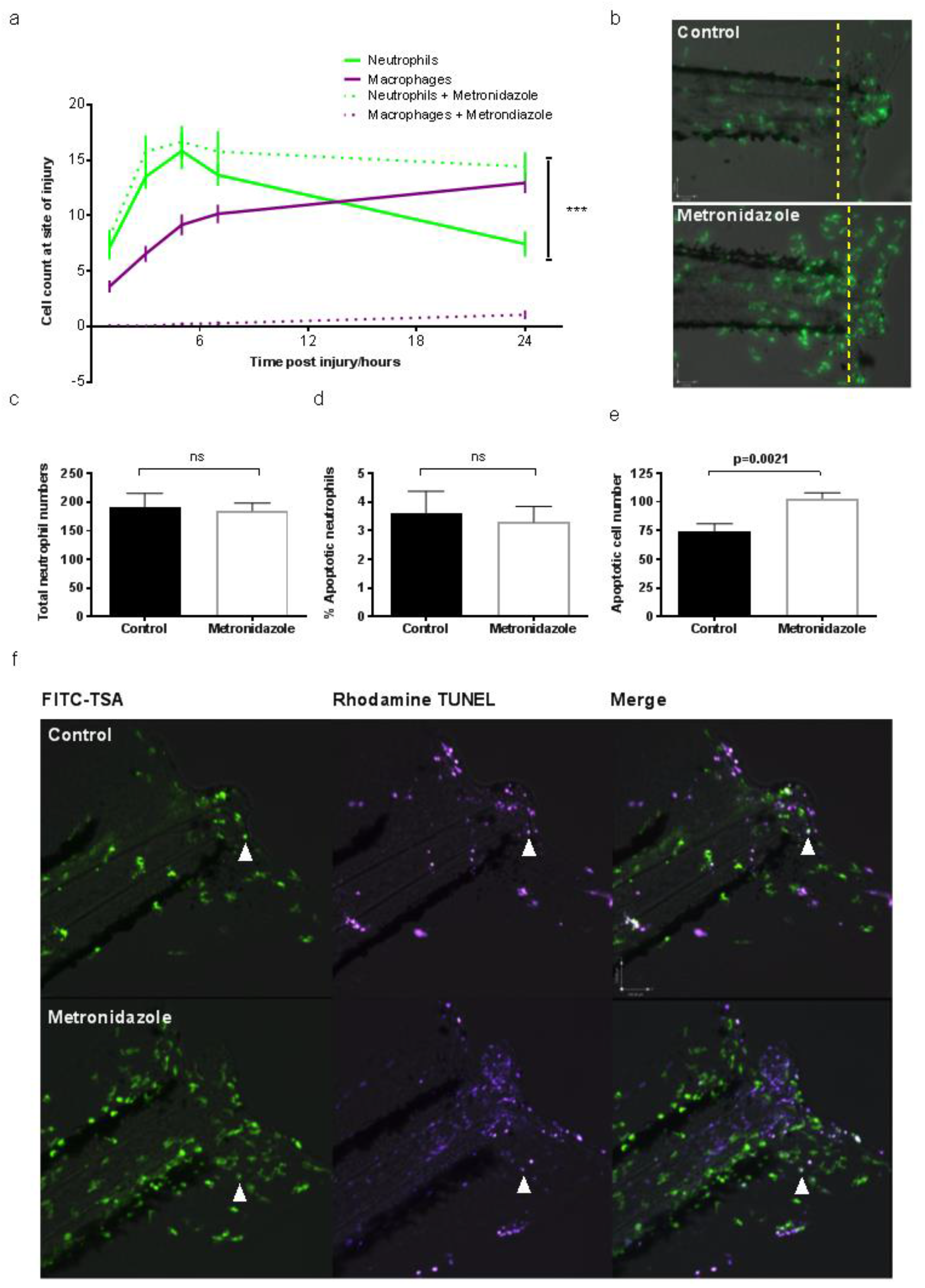
Macrophage clearance of apoptotic cells is necessary for successful resolution of neutrophilic inflammation in an *in vivo* zebrafish model. **(A)** Neutrophil and macrophage counts at the wound in triple transgenic *Tg(cfms:Gal4)i186;Tg(UAS:nfsB-mCherry)i149;Tg(mpx:EGFP)i114* larvae in the presence or absence of metronidazole. Neutrophil numbers are significantly higher at the wound in the absence of macrophages 24 hours post injury (hpi), p<0.0001. Statistics: t-test comparing NΦ counts 24 hours post injury with or without MΦ present. NΦ = Neutrophils. MΦ = Macrophages. All data are presented as mean +/− SEM, n=3, 6 fish per experiment. **(B)** Representative photomicrographs at 24 hpi, showing increased neutrophil numbers at the site of injury in metronidazole-treated larvae compared to control. Wound area classed as area to the right of the yellow dotted line. Images taken using 10x magnification on a TE2000U inverted microscope (Nikon). **(C)** Total neutrophil numbers are not affected in the absence of macrophages. **(D)** Dual FITC-TSA and Rhodamine-TUNEL positive cell counts in 8 dpf larvae 24 hpi in the presence or absence of metronidazole shows no significant difference (ns) in the percentage of apoptotic neutrophils. **(E)** Total apoptotic cell counts of TUNEL positive cells in 8 dpf larvae 24hpi. Macrophage-depleted larvae have significantly more apoptotic bodies at the wound, p=0.0021. All data are presented as mean +/− SEM, n=2, 6 fish per experiment for **(D)** and **(E)**. **(F)** Representative photomicrographs of 8 dpf larvae, 24 hpi treated with either DMSO control or metronidazole, and dual stained with FITC-TSA to label neutrophils and Rhodamine-TUNEL to label apoptotic cells at the wound (white arrow head). Images taken using 10x magnification on a TE2000U inverted microscope (Nikon).

In many models of inflammation, including the zebrafish, apoptosis of neutrophils contributes to inflammation resolution (*6*, *31*). It might therefore be expected that macrophage depletion would leave apoptotic neutrophils at the wound site. To test whether the persistence of apoptotic corpses explained the increase in neutrophil numbers, we ablated macrophages by treating transgenic zebrafish with metronidazole at 7 dpf, injuring the tailfin as before at 8 dpf and then fixing at 24 hpi in 4% paraformaldehyde for TUNEL staining. Older larvae were used to increase cell numbers involved in the inflammatory response and to increase our ability to detect apoptotic events. We co-stained for endogenous peroxidase activity of neutrophils using a modification of the Tyramide Signal Amplification system (*6*). Double stained cells corresponding to apoptotic neutrophils were then counted in macrophage-depleted larvae. Although the number of neutrophils was higher in the metronidazole treated fish, the percentage of these neutrophils that were apoptotic was not different in the metronidazole treated group (Figure 1d), suggesting that persisting apoptotic neutrophil corpses did not account for the increase in neutrophil numbers in the absence of macrophages. During analysis of these images, we did notice a striking and unexpected increase in TUNEL positive, TSA negative cells in macrophage-depleted larvae (Figure 1e), corresponding to apoptotic cells from non-neutrophil lineages. Representative images of stained cells are shown in Figure 1f. This demonstrates that macrophages have an important role in removing apoptotic cells arising from tissue injury and that neutrophil persistence is not due to un-cleared apoptotic neutrophils. This suggested to us that the majority of neutrophils might be removed from the wound by an alternative mechanism, such as reverse migration (*12*).

### Inflammation resolution is not exclusively dictated by direct macrophage-neutrophil interaction

Neutrophil-macrophage interaction has been observed during the inflammatory response (*32*) and under some circumstances, this has been shown to alter neutrophil migratory behaviour in ways that could explain the observed failure of inflammation resolution in the absence of macrophages. We therefore sought to analyse neutrophil migration before and after interaction with macrophages to see whether that interaction caused neutrophils to migrate away from the wound, thereby dissipating the inflammatory burden. We tracked individual transgenically-labelled macrophages and neutrophils cells, monitoring their interaction with other inflammatory cells. Neutrophils and macrophages were tracked during an inflammatory response and their x,y coordinates plotted (Figure 2a). Interaction between red (macrophage) and green (neutrophil) tracks was identified, and for each interaction we identified whether contact between a macrophage and neutrophil induced a significant change in behaviour of the neutrophil. Direct cell contact was defined as when membranes of both cell types could be seen to touch. Interaction with a macrophage made no significant difference in the neutrophil meandering index (Figure 2b), suggesting that interaction did not lead to a change in migratory behaviour. Moreover, analysis of vector maps of neutrophils pre- and post-interaction with a macrophage showed no clear difference between the 2 vector maps. The vector maps show the overall direction a neutrophil takes during the analysed period (Figure 2c). When analysed, the percentage of reverse migrated neutrophils that did not interact with a macrophage was 71%, compared to 65% of neutrophils reverse migrating following a macrophage interaction (Figure 2d). There was no significant difference in reverse migration behaviour following macrophage interaction (Fisher’s exact test p=0.78). Direct contact between macrophages and neutrophils is therefore not sufficient to completely explain the observed changes in neutrophil direction, suggesting an additional soluble factor is involved.

**Figure 2.**
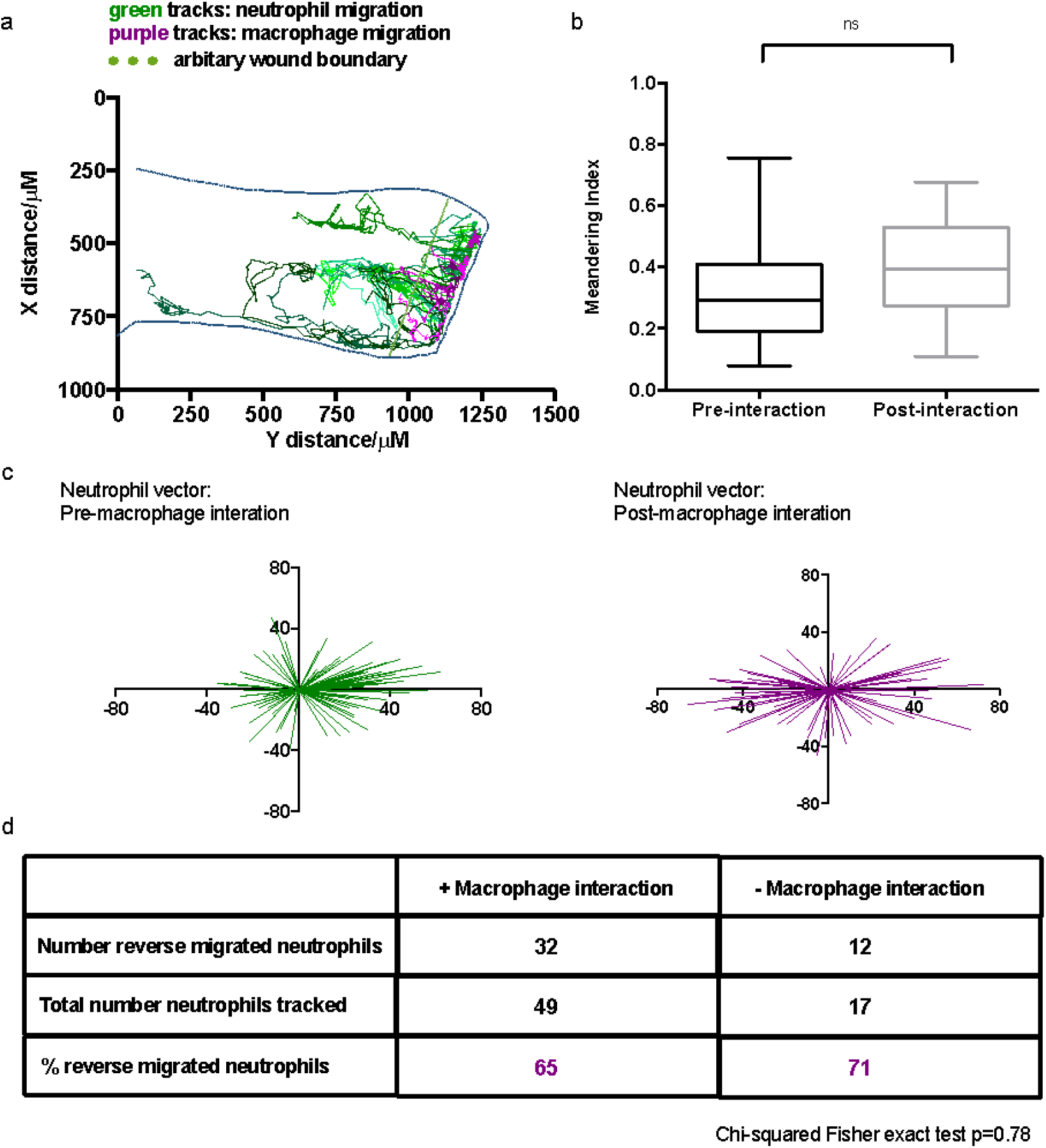
Inflammation resolution is not exclusively dictated by direct macrophage-neutrophil interaction. **(A)** Neutrophil (8 green lines) and macrophage (6 magenta lines) tracks during an inflammatory response are plotted using their x,y coordinates. Larval caudal fin outline superimposed over tracks to indicate position at wound. **(B)** The meandering index (migration pattern) of neutrophils before and after contact with a macrophage shows no significant difference. Number of cells tracked: 14. Two-tailed paired t-test. **(C)** Representative vector maps of overall neutrophil direction pre-(green) and post-macrophage (magenta) interaction. **(D)** 65% of neutrophils that interacted with a macrophage reverse migrated, whereas 71% neutrophils without a macrophage interaction also reverse migrated away from the wound site. Chi-squared Fisher exact test indicated no significant difference in reverse migration with or without macrophage contact. *P*=0.78.

### The eicosanoid PGE_2_ is necessary for timely inflammation resolution

Previous studies *in vitro* have shown that, following uptake of apoptotic cells, macrophages can alter both their profile of cytokine production and their phenotype, from pro-inflammatory to anti-inflammatory (*21*). It has been reported that levels of TGFβ1 and PGE_2_ increase following phagocytosis of apoptotic cells, whereas levels of cytokines such as IL-10 and TNFα fail to increase further. When PGE_2_ production was blocked by indomethacin, TGFβ1 levels remained low, suggesting PGE_2_ might be an upstream regulator of macrophage phenotype following uptake of apoptotic cells (*21*). We therefore sought to establish whether PGE_2_ could be a potential mediator in enhancing inflammation resolution following tail injury *in vivo*.

Microsomal prostaglandin E synthase, Ptges, is necessary for the conversion of prostaglandin H_2_ into PGE_2_ (*33*). PGE_2_ levels can be significantly reduced when Ptges expression is inhibited through the use of a morpholino antisense oligo (*34*). We used a previously published morpholino to knockdown *ptges* to test the effect of reduced PGE_2_ levels on neutrophil behaviour during inflammation. Reverse Transcriptase PCR demonstrated a significant reduction in correctly spliced *ptges* transcript at 4 dpf (Supplemental data 1). Neutrophil counts throughout the inflammatory response revealed a significant increase in neutrophil recruitment to the wound site. At the later time points, during the resolution phase, significantly more neutrophils were retained at the site of injury (Figure 3a). Since PGE_2_ has been shown to be critical in haematopoiesis (*35*), we looked at total neutrophil number in *ptges* morphants. Knockdown of *ptges* significantly reduced the number of neutrophils in the whole zebrafish (Figure 3b), demonstrating the increase in neutrophil numbers seen during inflammation is not due to an overall increase in neutrophils within the zebrafish. These data demonstrate Ptges activity plays an important role in the resolution of inflammation.

**Figure 3.**
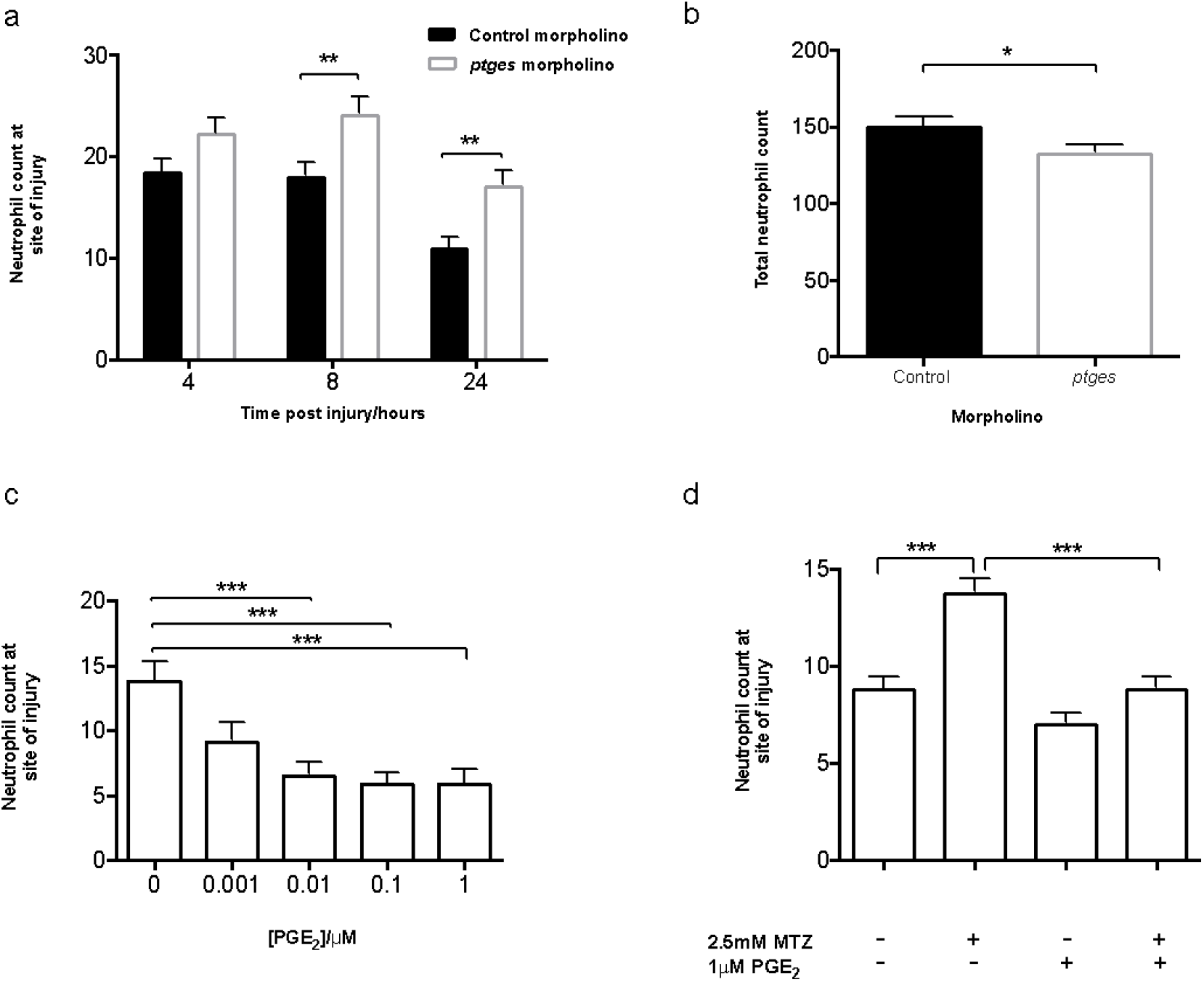
*In vivo* effects of PGE_2_ on neutrophils during an inflammatory response. **(A)** Morpholino knockdown of endogenous *ptges* shows the necessity of PGE_2_ during inflammation resolution, with a significant increase in neutrophil numbers at the wound site at 8 and 24hpi. *P*<0.01. **(B)** Total neutrophil numbers in unstimulated larvae are reduced (P<0.05) in the absence of *ptges*, indicating the increase during an inflammatory response is not due to increased neutrophils numbers overall. **(C)** Dose response showing increasing concentrations of PGE_2_ significantly drives neutrophilic inflammation resolution. PGE_2_ was added at 8 hpi with neutrophil counts performed at 12 hpi. Neutrophil numbers at the site of injury are significantly reduced between 0.01 μM and 1 μM. **(D)** In the absence of macrophages, exogenous PGE_2_ is able to significantly reduce neutrophil numbers back to basal levels at 24hpi. ***= *P*<0.001. One-way ANOVA, Bonferroni post-test. n=3, minimum of 14 larvae per experiment.

### PGE_2_ is sufficient to drive inflammation resolution

PGE_2_ has both anti- (*36*)and pro-inflammatory effects dependent on the timing of its production and concentration (*24*, *37*). To test whether PGE_2_ is sufficient to drive inflammation resolution, we immersed 3 dpf transgenic zebrafish larvae in a range of PGE_2_ concentrations at 8 hours after tailfin transection. Wound neutrophil numbers were then assessed at 12 hpi to look for accelerated inflammation resolution. Neutrophil numbers were significantly decreased in a dose-dependent manner when treated with PGE_2_, with 1 μM chosen as the optimal concentration (Figure 3c). Exogenous PGE_2_ was able to rapidly enhance neutrophil removal from the site of injury to return neutrophil numbers to basal levels, thereby promoting successful inflammation resolution. As PGE_2_ reduced neutrophil numbers at the site of injury, we tested if PGE_2_ accelerated neutrophil apoptosis at the wound site. 3 dpf larvae were injured, treated with PGE_2_ and fixed at 12 hpi for dual TUNEL (apoptosis) and TSA (neutrophil) staining. There was no difference in apoptotic cell numbers in larvae treated with PGE_2_ compared to control (Supplemental data 2a.). This suggests that neutrophils are removed through a mechanism other than apoptosis.

Since exogenous PGE_2_ could enhance inflammation resolution in our model, we investigated whether PGE_2_ could rescue the persisting neutrophil phenotype observed with macrophage ablation. Zebrafish larvae with Nitroreductase-expressing macrophages and GFP expressing neutrophils were treated with metronidazole at 2 dpf, injured at 3 dpf and treated with PGE_2_ at 8 hpi. Cell counts were performed at 12 and 24 hpi. PGE_2_ was able to significantly reduce neutrophil numbers at the wound in metronidazole-treated larvae at 12 hpi compared to metronidazole only larvae, with means of 4.6+/−0.4 and 9.6+/−0.8 respectively (Supplemental data 2b). At 24 hpi, neutrophil numbers remained significantly reduced (Figure 3d). These data demonstrate that PGE_2_ can drive inflammation resolution in the absence of macrophages, thus correcting the resulting inflammatory phenotype.

### Neutrophil migration pattern, but not speed, is modulated by PGE_2_

During inflammation resolution, neutrophils can be removed through multiple mechanisms, including neutrophil apoptosis (*38*) and reverse migration (*13*, *39*). Macrophage-depletion did not result in a detectable difference in apoptotic neutrophil numbers compared to control, due to either residual macrophage activity or the infrequency of neutrophil apoptosis in the zebrafish model. Nonetheless, PGE_2_ can still reduce neutrophil numbers in these larvae. This implies that neutrophils involved in the inflammatory response are being removed from the site of injury through a mechanism other than efferocytosis. We therefore investigated whether PGE_2_ could enhance inflammation resolution by acceleration of neutrophil reverse migration in tail-transected zebrafish larvae. Using well established photoconversion protocols (*40*, *41*) in transgenic zebrafish expressing the photoconvertable fluorophore *kaede* in neutrophils, we labelled neutrophils at the wound site at 4 hours after tailfin transection and observed the effects of addition of PGE_2_. Neutrophils at the wound site were tracked during the resolution phase of inflammation for 2 hours, from 10-12 hpi. Red, photoconverted neutrophils could be seen migrating away from the site of injury more readily in PGE_2_ treated larvae (Figure 4). PGE_2_ significantly increased the number of neutrophils moving away from the wound area (Figure 4d). These data show that PGE_2_ drives inflammation resolution by enhanced reverse migration away from the injury site.

**Figure 4.**
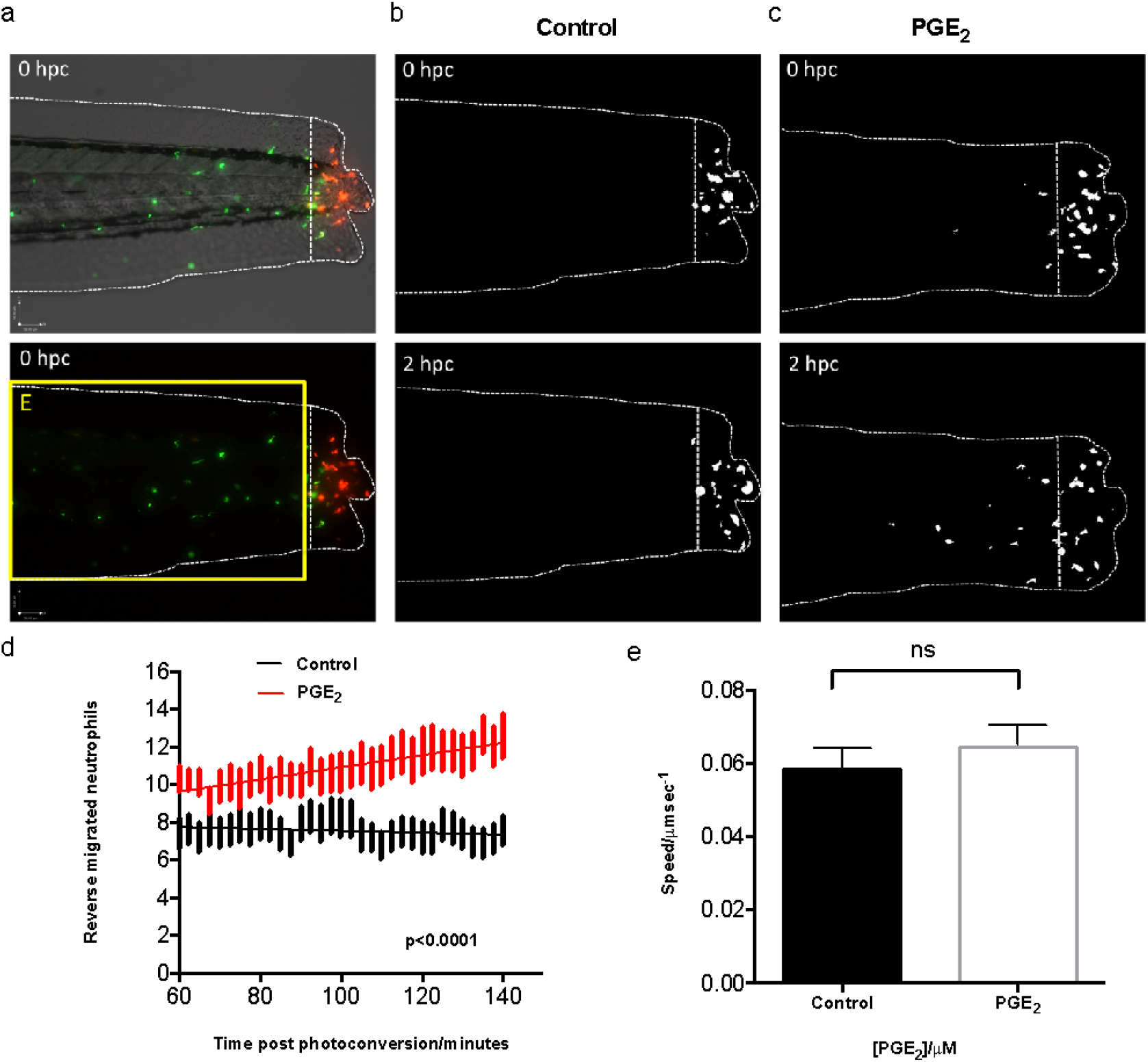
PGE_2_ drives inflammation resolution through accelerated reverse migration. **(A)** Representative photomicrographs of control or PGE_2_-treated 3dpf *Tg(mpx:gal4;UAS:kaede)i222* larvae following photoconversion. The area to the right of the white dotted vertical line indicates wound site, where cells were photoconverted from green to red fluorescence. The yellow box indicates the area into which the photoconverted cells migrate and corresponds to data in panel (d). **(B)** The red channel only is shown as a binary image, of a control larva at 0 and 2 hours post conversion (hpc). Very little migration away from the wound site occurs between 10 and 12 hours post injury **(hpi)**. **(C)** Binary images of the red channel of a PGE_2_-treated larva, at 0 and 2 hpc. At 12hpi, neutrophils have migrated away from the wound site, when treated with PGE_2_. **(D)** Plot showing the number of neutrophils moving away from the wound over 10-12 hpi, pre-incubated with or without PGE_2_ from 8-9 hpi. PGE_2_-treated neutrophils migrate further away from the site of injury between 10-12 hpi. Line of best fit shown is calculated by linear regression. *P* value shown is for the difference between the 2 slopes. **(E)** The speed of neutrophils moving away from the site of injury is not significantly different in the presence of exogenous PGE_2_, indicating neutrophils migrate away sooner rather than at a greater speed in PGE_2_-treated larvae. All data are presented as mean +/− SEM, n=3, 6 fish per experiment. Images taken using 10x magnification on a TE2000U inverted microscope (Nikon).

We then asked whether this enhanced migration from the wound could be due to an increased migration speed of the neutrophils or to them leaving the site of injury sooner. Neutrophil speed away from the site of injury was measured in the same reverse migration assays described above (Figure 4e). There was no significant difference in migration speed, implying that PGE_2_ may enhance release of neutrophils from their patrolling behaviour at the wound site. The timing of PGE_2_ release is therefore critical to allow neutrophils to perform their role and then drive them away without compromising the inflammatory response.

### Signalling through EP4 receptors contributes to inflammation resolution

PGE_2_ can signal through 4 prostanoid receptors: EP1, EP2, EP3 and EP4 (*42*). The expression and distribution of these receptors vary between different tissues and cell types. EP4 is considered a stimulatory receptor and is expressed on human macrophages, with studies showing that blockage of EP4 signalling inhibits cytokine release by macrophages (*43*). All 4 EP receptors are present in zebrafish, with multiple paralogues of each receptor (*44*). EP4b is the most abundant in adult zebrafish and is closest to human EP4 phylogenetically (*45*). To test whether PGE_2_ could be signalling *in vivo* through EP receptors we used a previously published antagonist, AH23848, to inhibit EP4 receptor signalling (*46*). Neutrophil counts at 24 hpi showed a significant increase in number when EP4 signalling was blocked, suggesting endogenous PGE_2_ may act via this receptor. Addition of PGE_2_ at standard doses did not lead to significant abrogation of this effect (Figure 5a). These data suggest PGE_2_ signals predominantly through the EP4 receptor, contributing significantly to inflammation resolution. However, it remained unclear how EP4 signalling might lead to altered neutrophil behaviour.

**Figure 5.**
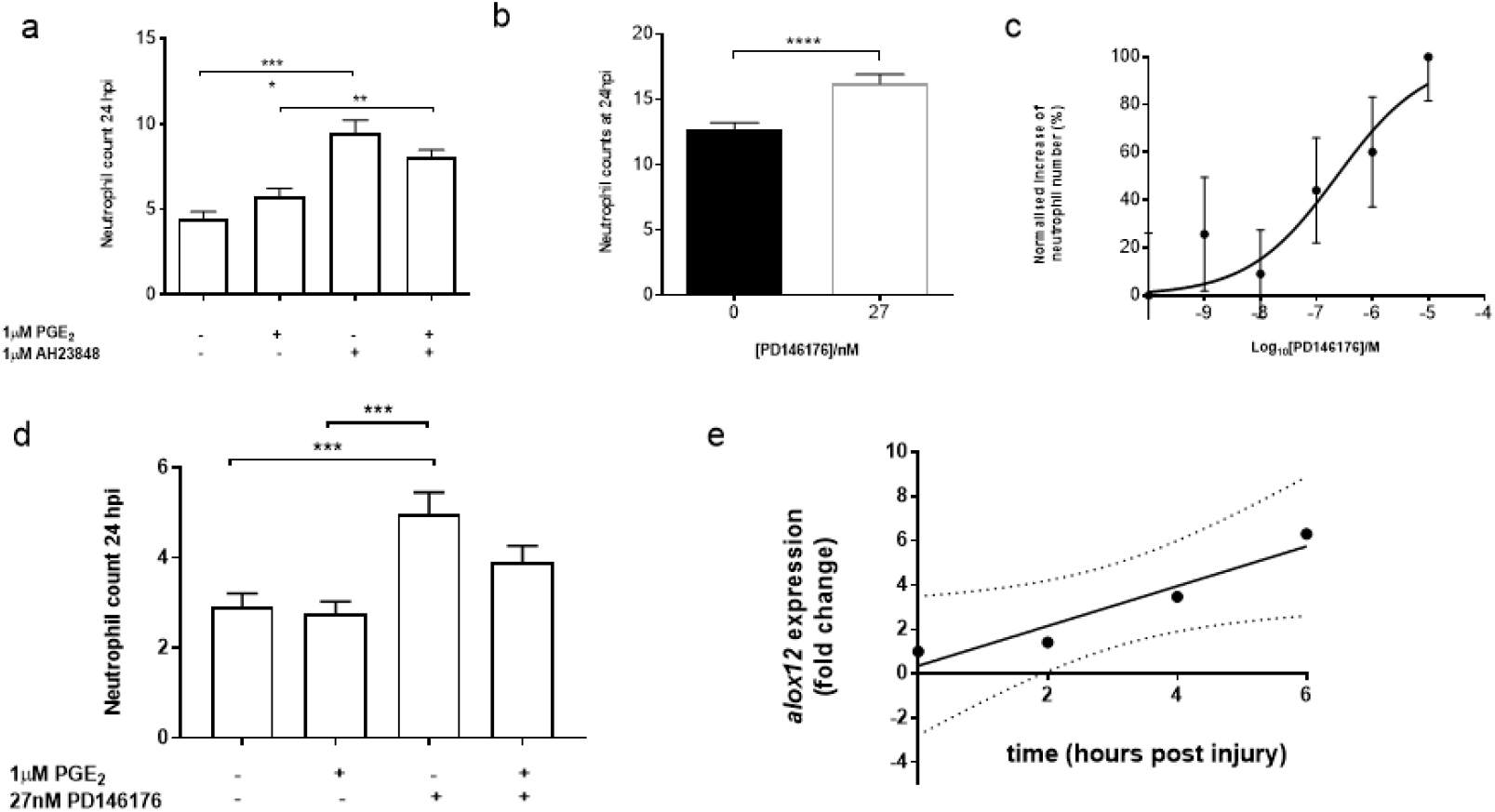
EP receptor signalling and downstream activation of lipoxygenases by PGE_2_ leads to lipid-mediator class switching and inflammation resolution. **(A)** Neutrophil counts at 24 hpi show a significant increase in neutrophil number when EP4 signalling is blocked using the antagonist AH23848. Addition of PGE_2_ does not lead to significant abrogation of this effect implying PGE_2_ signals through the EP4 receptor to promote neutrophil removal. **(B)** Downstream activity of 15-LO was inhibited using the antagonist PD146176, also causing a significant delay in neutrophil removal from the wound site at 24hpi. \*\*\**P*<0.001. Data represent 3 replicates with a minimum of 30 fish per replicate. T-test performed. **(C)** Varying doses of PD146176 increases neutrophil numbers at the wound site 24 hpi in a dose-dependent manner. **(D)** Exogenous PGE_2_ is unable to significantly decrease neutrophil numbers in the presence of the 15-LO inhibitor, PD146176, at 24hpi. **(E)** qPCR expression analysis of *alox12* following injury shows a significant increase up to the peak of inflammation. Fold change of expression to the reference gene represents the mean ± SEM of 4 replicate experiments per time point, with a minimum of 50 larvae per replicate. p=0.04, r^2^=0.91.

### Upregulation of lipoxygenase by PGE_2_ is necessary for inflammation resolution

The metabolism of the membrane lipid arachidonic acid is a complex pathway, producing not only prostaglandins, but also other eicosanoids such as pro-resolving lipoxins and pro-inflammatory leukotrienes. Lipoxygenases (LOXs) are essential for the production of these metabolites, with select isozymes being key for particular pathways. Previous studies have shown that arachidonate 12-lipoxygenase (12-LO) or 15-lipoxygenase (15-LO) activity, in combination with arachidonate 5-lipoxygenase (5-LO), can produce lipoxins, and 5-LO activity alone forms leukotrienes (*47*). Increasing concentrations of PGE_2_ inhibit 5-LO translocation from the cytoplasm to the nucleus, abrogating leukotriene synthesis (*48*), and skewing the balance between pro- and anti-inflammatory mediators towards increased lipoxin synthesis - a process termed “lipid-mediator class switching” (*28*). Zebrafish have the three key lipoxygenase genes for lipid mediator synthesis: *alox12, alox5* and *alox15b* genes (*49–51*). To assess the contribution of lipoxin production during inflammation resolution in the zebrafish, we immersed injured transgenic larvae in PD146176, a specific 15-LO inhibitor (*52*). We found that inhibiting 15-LO activity during inflammation led to a significant increase in neutrophil numbers at 24 hpi compared to control larvae (Figure 5b), indicating persisting inflammation at the wound site. This effect was dose dependant (Figure 5c). Addition of exogenous PGE_2_ following 15-LO inhibition, was unable to abrogate this effect, with neutrophil numbers remaining high at the wound site 24hpi (Figure 5d), supporting a mechanism of PGE_2_ acting upstream of 15-LO in this pathway.

The oxygenation of arachidonic acid by lipoxygenases can occur at varying positions along the fatty acid carbon chain (*47*), and can determine the functionality of the protein according to which carbon is oxygenated and which amino acid residue is present at specific sites. In zebrafish, LOX genes are annotated in Ensembl as *alox12* and *alox15b*, yet the presence of phenylalanine (F) at position 353 and a bulky valine (V) in position 418^1^ suggests that zebrafish *alox12* should function predominantly in the role of a 15-LO (Supplementary figure 3). We therefore assessed alteration of expression of *alox12* (the zebrafish putative 15-LO) by qPCR during injury-induced inflammation. There was a significant linear increase in 12-lipoxygenase expression up to 6 hours after injury (Figure 5e), demonstrating that inflammation caused by injury induces a change in lipoxygenase activity. This suggests a plausible mechanism for the induction of reverse migration through increased lipoxin production during inflammation resolution.

### Lipoxin A_4_ drives inflammation resolution via reverse migration

LipoxinA_4_ (LXA_4_) is a pro-resolving product of arachidonic acid metabolism that inhibits neutrophil chemotaxis, transmigration, superoxide generation and the production of pro-inflammatory cytokines (*53*). Because *alox12* expression was increased during injury-induced inflammation, we next determined if its key product LXA_4_ plays a role in inflammation resolution in the zebrafish model. First, we added exogenous LXA_4_ to the injured larvae and assessed neutrophil number at 6 hpi. Our data demonstrate a protective effect of LXA_4_, acting to inhibit neutrophil chemotaxis, significantly reducing neutrophil recruitment to the wound site (Figure 6a). If PGE_2_ production following injury increases *alox12* activity during inflammation resolution, and thereby increases LXA_4_ production, we would predict LXA_4_ could allow removal of neutrophils already present at the wound site due to loss of sensitivity to local wound signals. To assess this, we performed reverse migration experiments in the presence of exogenous LXA_4_. LXA_4_ caused neutrophils to move away from the wound earlier than in untreated larvae (Figure 6b), without altering neutrophil migration speed (Figure 6c). Our data are best explained by that production of LXA_4_ by Alox12 following injury-induced PGE_2_ production as the mechanism that promotes inflammation resolution *in vivo*.

**Figure 6.**
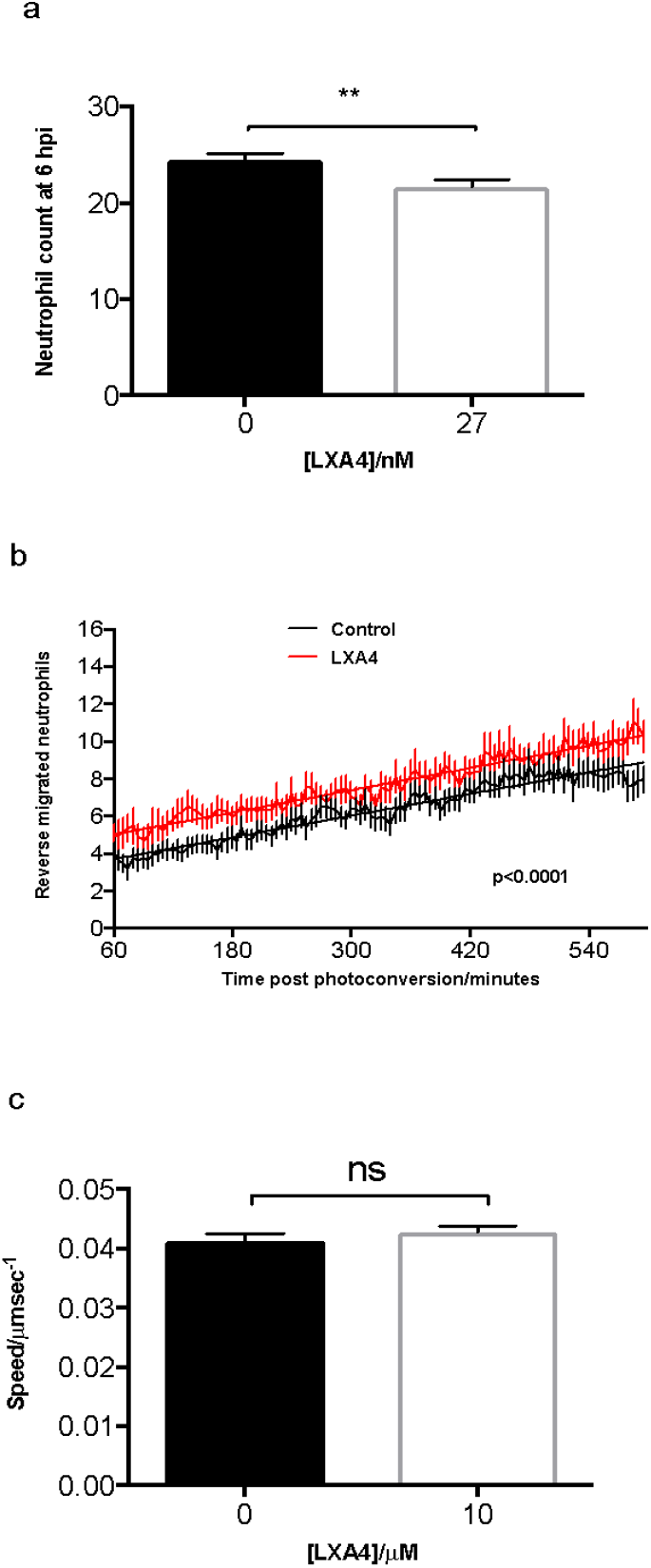
Lipoxin A_4_ drives inflammation resolution via reverse migration. **(A)** Injection of LXA_4_ into zebrafish larvae causes a small but significant reduction in neutrophil numbers at the wound site at 6 hpi. **(B)** Plot showing the number of neutrophils reverse migrating away from the wound over a time course of 6-15 hpi, pre-injected at 4 hpi with or without LXA_4_. LXA_4_-treated neutrophils migrate away from the site of injury sooner than control neutrophils. Line of best fit shown is calculated by linear regression. *P* value shown is for the difference between the 2 slopes. **(C)** The speed of neutrophils moving away from the site of injury was measured and is not significantly different in the presence of exogenous LXA_4_. All data are presented as mean +/− SEM, n=3, minimum 8 fish per experiment.

## Discussion

Restoration of tissue homeostasis following infection or injury requires successful resolution of inflammation. Inflammation resolution should not occur before the threat is ablated, nor should inflammation persist longer than necessary. The process is therefore tightly regulated. Neutrophils and macrophages as a population are well positioned to assess the threat to the organism and determine the duration of the inflammatory response. Once resolution is decided, neutrophils can either die at inflammatory sites by apoptosis and be cleared by macrophages (*54*) or can be removed from inflammatory sites by altered migratory patterns (reverse migration) (*39*). If this tightly controlled inflammatory process is disrupted, failure of inflammation resolution can lead to the tissue damage seen in chronic inflammatory diseases (*55*). The molecular mechanisms controlling the decisions on duration of inflammation are complex and yet to be fully determined. Existing evidence indicates that lipid mediator secretion is likely to be important in signalling inflammation resolution (*56*) but it is unknown which mediators regulate neutrophil removal from sites of inflammation *in vivo*. We have shown for the first time that the lipid mediator signalling molecule PGE_2_ is essential for determining the outcome of neutrophil inflammation in response to tissue injury. Moreover, our data suggest that alterations in Alox activity leads to production of LXA_4_. This lipid mediator class switching appears to be the critical mechanistic step by which macrophage PGE_2_ release after uptake of apoptotic cells promotes inflammation resolution *in vivo*.

Previous *in vitro* studies have suggested that phagocytosis of apoptotic neutrophils by macrophages might contribute towards successful inflammation resolution (*21*, *57*). Ingestion of apoptotic cells by macrophages promotes an anti-inflammatory environment, suppressing pro-inflammatory cytokine production and promoting LXA_4_ production (*21*). While mice lacking macrophages and neutrophils show healing without excessive inflammation or scar formation, in macrophage-only depleted mice, neutrophils populated the wound area, with impaired wound morphology and delayed healing (*58*). We have shown that in the absence of macrophages, apoptotic bodies accumulate at the wound site, resulting in neutrophil persistence and prolonged inflammation. This demonstrates the importance of understanding macrophage-neutrophil interactions during inflammation resolution to prevent further damage to healing tissues.

Recent work has shown that reverse migration is a contributor to neutrophil removal from inflammatory sites (*13*, *15*, *39*, *59*). It has been shown that macrophage recruitment and redox-SFK (Src family kinase) signalling plays a key role in neutrophil-mediated inflammation resolution (*32*). These studies also suggest a direct contact mechanism between neutrophils and macrophages, which may direct neutrophil reverse migration; however, in the absence of macrophage contact, neutrophils are still able to reverse migrate, suggesting that other, unknown factors may be involved. Tracking analysis of macrophage and neutrophil contact in our model of inflammation demonstrates that direct contact is not an essential requirement for reverse migration. Furthermore, our studies support the importance of a soluble factor, the lipid mediator PGE_2_ for successful inflammation resolution. Our data suggest that this lipid mediator is a key signalling molecule between macrophages and neutrophils during inflammation, leading to production of LXA_4_, enhancing neutrophil reverse migration and hence speeding inflammation resolution.

PGE_2_ is known to be involved in many cellular processes, including cell proliferation, apoptosis, angiogenesis, inflammation and immune surveillance (*60*). The production of PGE_2_ from prostaglandin synthases is essential for development of organisms past the gastrulation stage (*34*) and is crucial for hematopoietic stem cell homeostasis (*35*). However, to our knowledge, PGE_2_ has not been shown to be necessary for resolution of neutrophilic inflammation. We have shown by genetic and pharmacological approaches that the production of PGE_2_ is essential for inflammation to resolve, and that PGE_2_ is sufficient to drive neutrophil removal from wound sites. PGE_2_ can promote neutrophil removal from wound sites in the absence of macrophages, implying an important connection between macrophages and the lipid mediator PGE_2_. This could prove to be a useful target for therapeutic intervention in chronic inflammatory disease.

PGE_2_ acts via EP receptors present on a range of cell types, including neutrophils, macrophages, epithelial and endothelial cells. Neutrophils predominantly express EP2 and EP4 (*61*), both of which, upon activation, cause an increase in cAMP levels within the cell. An increase in intracellular cAMP can “reprogram” the cell to switch from 5-LO products to the production of 15-LO products (*28*). Our studies show PGE_2_ signalling through EP4 receptors to be important in the resolution of neutrophilic inflammation and that this activity is mediated via 15-LO products, most likely the anti-inflammatory mediator LXA_4_.

Human PMNs *in vitro* dramatically increase expression of 15-LO mRNA in the presence of PGE_2_ (*28*). Our data from whole animals shows a significant increased expression of the annotated zebrafish 12-LO mRNA following tailfin injury relative to time. Human blood neutrophils exposed to PGE_2_ inhibit LTB_4_ biosynthesis, and demonstrate reduced neutrophil migration (*62*). Our data confirm the regulation of neutrophil migration by PGE_2_ and LXA_4_ in an *in vivo* model of inflammation resolution. Neutrophils more readily migrate away from the wound site when exposed to exogenous PGE_2_ or LXA_4_. The addition of PGE_2_ after the maximum recruitment of neutrophils to the wound is able to dramatically enhance the reverse migration of neutrophils. The mechanism of this step is still unknown, but PGE_2_ alteration of LO activity would fit with its known role in inhibiting neutrophil recruitment (*63*). In our studies, exogenous LXA_4_ causes neutrophils to leave the wound sooner. LXA_4_ has been shown to inhibit chemokine signalling in neutrophils *in vitro* (*64*), suggesting a mechanism by which LXA_4_ facilitates neutrophil reverse migration from sites of inflammation. Mathematical modelling data from *in vivo* neutrophil migration following zebrafish tail fin wounding suggests that neutrophil reverse migration is a phenomenon of stochastic redistribution of neutrophils in the tissues (*65*). This suggests a model whereby neutrophils are retained at wounds by a molecular “retention signal” and that decay of this signal leads to reverse migration (*12*). Chemokine receptor desensitisation would be strong candidates for explaining how these retention signals might be initiated and subsequently decayed. LXA_4_ interference with these signals might be a potential mechanism by which PGE_2_ ultimately alters neutrophil migration patterns leading to inflammation resolution.

In this context, an *in vivo* reporter would be useful to further elucidate the presence and activity of PGE_2_. Understanding the molecular mechanisms involved in neutrophil removal from wound sites, specifically during reverse migration, would greatly improve our understanding of inflammatory disease and provide potential strategies and targets for much needed therapeutic intervention.

## Material and methods

### Maintenance and breeding of zebrafish

To visualise both macrophages and neutrophils within the same zebrafish larva, triple transgenic fish *Tg(cfms:Gal4)i186;Tg(UAS:nfsB-mCherry)i149; Tg(mpx:EGFP)i114*, previously described (Gray, 2011) were incrossed, subsequently termed *fms:UNM:mpx. Tg(mpx:gal4; UASkaede)i222* zebrafish were used for photoconversion and neutrophil tracking experiments, subsequently termed *mpx:kaede. Tg(mpx:EGFP)i114* zebrafish were used to study neutrophils during inflammation. Zebrafish strains were maintained according to standard protocols. Adult fish were maintained on a 14:10-hour light/dark cycle at 28°C in UK Home Office approved facilities in the MRC Centre for Developmental and Biomedical Genetics aquaria at the University of Sheffield.

### Inflammation assay

Inflammation was induced in zebrafish embryos by tail transection as described previously (*66*). Embryos were anesthetized at 3 or 8 days post fertilization (dpf) by immersion in 0.168 mg/mL Tricaine (Sigma-Aldrich), and tail transection was performed using a microscalpel (World Precision Instruments). Neutrophils and macrophages were counted at the site of transection at various timepoints including 2, 4, 6, 8 and 24 hours post injury (hpi) using a fluorescent dissecting stereomicroscope (Leica).

### In vivo cell-specific ablation

Metronidazole (Sigma-Aldrich) was dissolved to a concentration of 2.5mM in E3 with 0.2% DMSO. Solutions of Metronidazole were made up fresh on the day of use with vigorous agitation until fully dissolved. *Tg(cfms:Gal4)i186;Tg(UAS:nfsB-mCherry)i149; Tg(mpx:EGFP)i114* were immersed in Metronidazole or E3 and 0.2% DMSO, at 2 or 7 dpf overnight at 28 degrees Celcius for 15 hours. Larvae were wrapped in foil during the experiment. At 3 or 8 dpf, larvae were checked for macrophage depletion on a fluorescent dissecting stereomicroscope (Leica).

### Neutrophil and apoptotic cell labelling

Rates of neutrophil apoptosis were assessed by dual staining with Rhodamine-TUNEL and FITC-Tyramide Signal Amplification (TSA). Transgenic zebrafish larvae were injured at 8 dpf and fixed at 24 hpi in 4% paraformaldehyde. TUNEL (ApopTag Red; Millipore Corporation) staining labelled apoptotic cells with red fluorescence, and TSA (TSAplus kit; Fluorescence Systems, PerkinElmer Life and Analytical Sciences) staining labelled neutrophils with fluorescein green fluorescence. Neutrophils at the wound were imaged on an Eclipse TE2000-U inverted compound fluorescence microscope (Nikon), and neutrophil specific apoptosis was assessed by the percentage of TSA-positive neutrophils labelled with TUNEL. Red TUNEL positive only cells were assessed for total apoptosis rates.

### Cell tracking

To assess neutrophil migration behaviour pre- and post-interaction with a macrophage, transgenic larvae *Tg(mpx:EGFP)i114* were crossed to *Tg(mpeg1:mcherry^caax^)SH378*. 3 dpf larvae were injured and images were taken every 3 minutes between 0 - 12 hpi using an Eclipse TE2000-U inverted compound fluorescence microscope (Nikon). Direct contact was defined as when cell membranes could be seen to touch. Comparisons of migration behaviour were made over a 30 minute period pre- and post-interaction.

### Compound treatment of zebrafish larvae

Following tail transection at 3 dpf, zebrafish larvae were screened at 4 hpi for neutrophil recruitment. Selected larvae were immersed in 1μM PGE_2_ (Sigma-Aldrich) at 8 hpi with counts performed at 12 hpi, to look for accelerated inflammation resolution and 24 hpi for correction of phenotype assays. Concentrations of PGE_2_ ranged from 0.01μM to 10μM for dose response assays. The EP4 antagonist AH23848 (Cayman Chemicals) was used at a final concentration of 1μM. Fish were incubated at 2 dpf and with tail transection being performed at 3 dpf. PD146176, a 15-LOX inhibitor (Sigma), or Lipoxin A_4_ (LXA_4_) (Cayman Chemicals) were injected into the Duct of Cuvier to a final concentration of 27nM at 2dpf, followed by tail transection and 6 or 24 hpi counts.

### Reverse migration assay

Tail transection of *Tg(mpx:gal4;UASkaede)i222* larvae was performed at 3 dpf. Embryos were raised to 8 hpi, incubated in 1μM PGE_2_ for 1 hour and mounted in 0.8% low-melting-point agarose (Sigma-Aldrich). An UltraVIEW PhotoKinesis device on an UltraVIEWVoX spinning disk confocal microscope (PerkinElmer Life and Analytical Sciences) was used to photoconvert Kaede–labelled cells at the wound site using 120 pulses of the 405 nm laser at 40% laser power. Embryos were transferred to an Eclipse TE2000-U inverted compound fluorescence microscope (Nikon), where a 1394 ORCA-ERA camera (Hamamatsu Photonics Inc) was used to capture a time-lapse series with 2.5 minute intervals for 2 hours. Tracking analysis of red fluorescing neutrophils was performed in Volocity 6 (Improvision; PerkinElmer Life and Analytical Sciences), using the intensity of fluorescence to identify individual labelled neutrophils over time. LXA_4_ was injected at 4 hpi at a final concentration of 27nM, neutrophils at the wound site photoconverted at 5 hpi and larvae were timelapsed between 6 and 15 hpi. Speed data were calculated using tracking timelapses where larvae were incubated in 10μM LXA_4_.

### Morpholino studies

All morpholinos were obtained from GeneTools LLC, Philomath, OR, USA. *ptges* splice blocking morpholino is described by Feng et al, 2012. 0.5 nl of 0.4mM of standard control morpholino (CoMo) or *ptges* morpholino were injected into embryos at the 1 cell stage previously described (Feng et al., 2012). Injected larvae were incubated in 5 μM PGE_2_ at the onset of epiboly to compensate for the PGE_2_ required for normal development up to 42 hpf. PGE_2_ was then withdrawn from the larvae at 48 hpf. At 3 dpf, larvae were injured and neutrophil counts performed at 4, 8 and 24 hpi. Knockdown was assessed in the same fish at 48 hpf and 4 dpf using Reverse Transcriptase PCR, primer sequences were PTGESF1 5’-CTCGGGAGCGACATACAGTT-3’ and PTGESR1 5’-AGCAGATATGCAACGCTGTG-3’. *ptges* splice blocking morpholino leads to deletion of exon 2 and a loss of 83 bp.

### qPCR

3 dpf larvae were injured and total RNA was extracted using Trizol at various time points post injury. cDNA synthesis was performed using Superscript II (Thermo Scientific) and qPCR experiments using DyNAmo Flash SYBR green qPCR kit (Thermo Scientific) were carried out. ΔCT was calculated using *elongation factor 1 alpha* as a reference gene. Relative expression levels were plotted after determining ΔΔCt by normalizing to a single sample with a high ΔCT value. Primer sequences for *alox12* gene were alox12F ccacggaatatcaccagatgga and alox12R attcagagtcgttgtgggtatc.

### Statistical analysis

Data were analysed (Prism 7.0, GraphPad Software, San Diego, CA, USA) using unpaired, two-tailed t-tests for comparisons between two groups and one-way ANOVA (with appropriate post-test adjustment) for other data. A Chi-squared Fisher’s exact test was used to assess statistical significance in Figure 2. Linear regression analysis was performed on qPCR data in figure 5.

## Acknowledgements

General: We are grateful for the use of the Bateson Centre aquarium at the University of Sheffield and their support. Funding: This work was supported by a Medical Research Council (MRC) Senior Clinical Fellowship with Fellowship-Partnership Award and MRC Programme Grant to S.A.R. (G0701932 and MR/M004864/1) and MRC Centre Grant G0700091. Funding for Y.F was provided by Wellcome Trust Sir Henry Dale Fellow 100104AIA. Funding for B.D.L is an NIH grant (P01GM095467).

## Author contributions

S.A.R planned the study and designed the experiments with contribution from C.A.L, F.E and B.D.L. C.A.L, J.A.L, A.L.R, M.J.G.S and F.E. performed the experiments. Y.F, B.D.L, and M.K.W provided scientific expert knowledge. C.A.L and S.A.R analyzed the data. C.A.L and S.A.R wrote the manuscript with assistance from all authors.

## Data and materials availability

Materials can be obtained through an MTA.

## Supplementary Materials

**Supplementary Figure 1.**
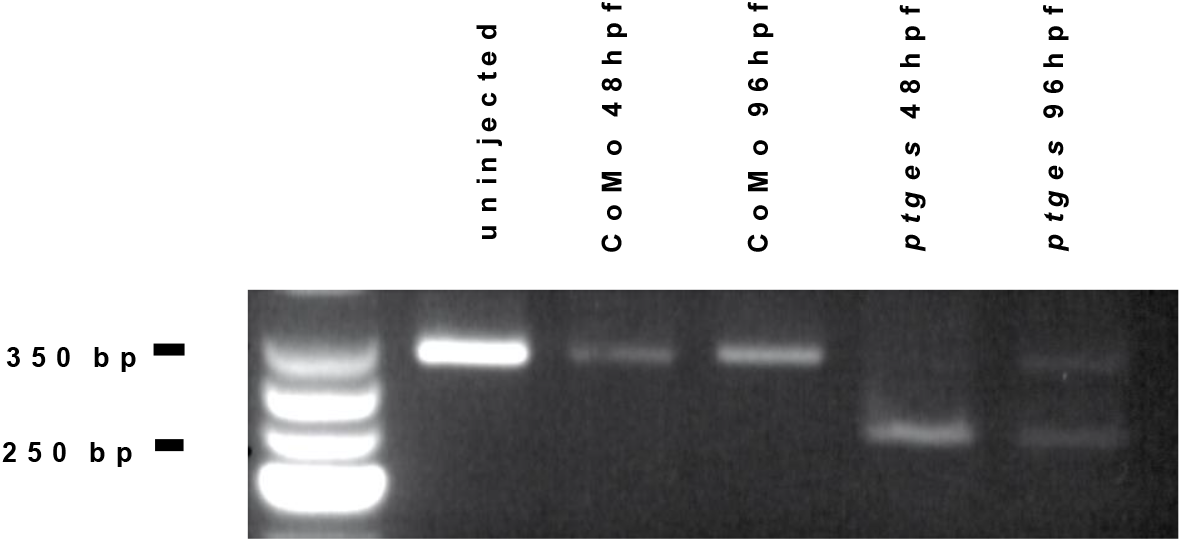
RT-PCR demonstrates successful exon deletion. Gel electrophoresis showing almost complete knockdown of *ptges* at 48 and partial knockdown remaining at 96 hpf. A splice variant missing exon 2 in *ptges* morphants of 250bp compared to a WT product of 350bp is visible.

**Supplementary Figure 2.**
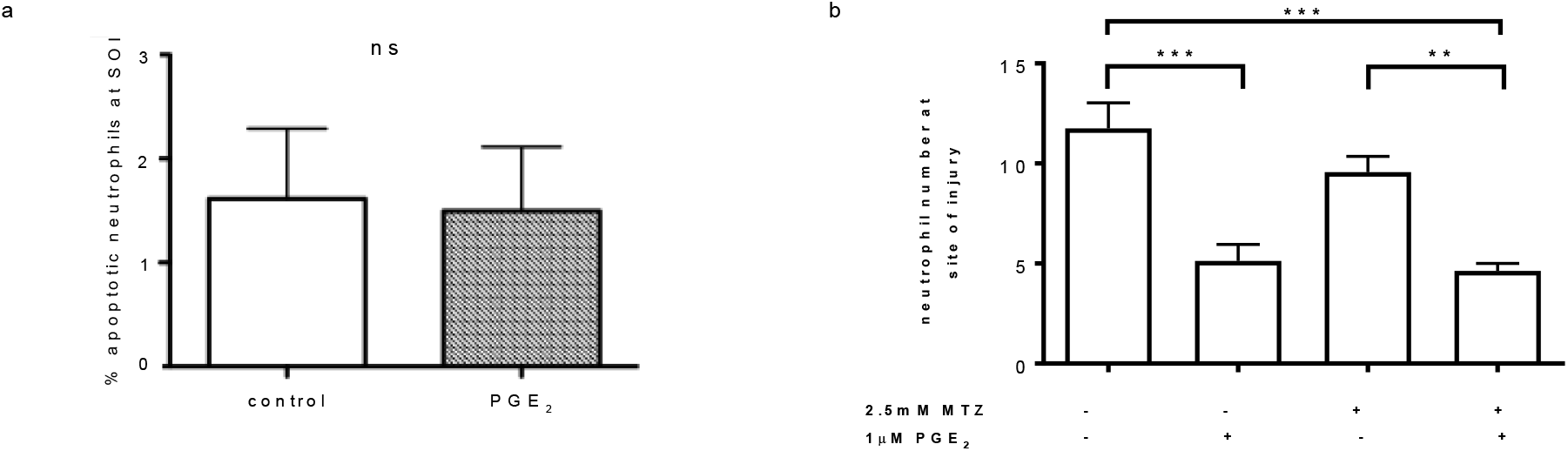
PGE_2_ drives resolution of inflammation in the absence of macrophages by 12 hpi and does not alter neutrophil apoptosis. **(A)** Graph showing PGE_2_ does not affect neutrophil apoptosis during inflammation resolution. Larvae were injured, treated with PGE_2_ and fixed at 12 hpi for TSA and TUNEL staining. The percentage of apoptotic neutrophils at the site of injury was not altered by the addition of PGE_2_. n=2, minimum of 18 fish per experiment. **(B)** Neutrophil counts 12 hpi in 3 dpf *Tg(mpx:EGFP)i114;Tg(cfms:Gal4)i186;Tg(UAS:nfsB-mCherry)i149* larvae treated with and without 2.5mM metronidazole and 1μM PGE_2_. PGE_2_, added at 8 hpi, can significantly drive inflammation resolution by 12 hpi, even when macrophages are still being recruited to the site of injury in control larvae or in their complete absence when treated with metronidazole.

**Supplementary Figure 3.**
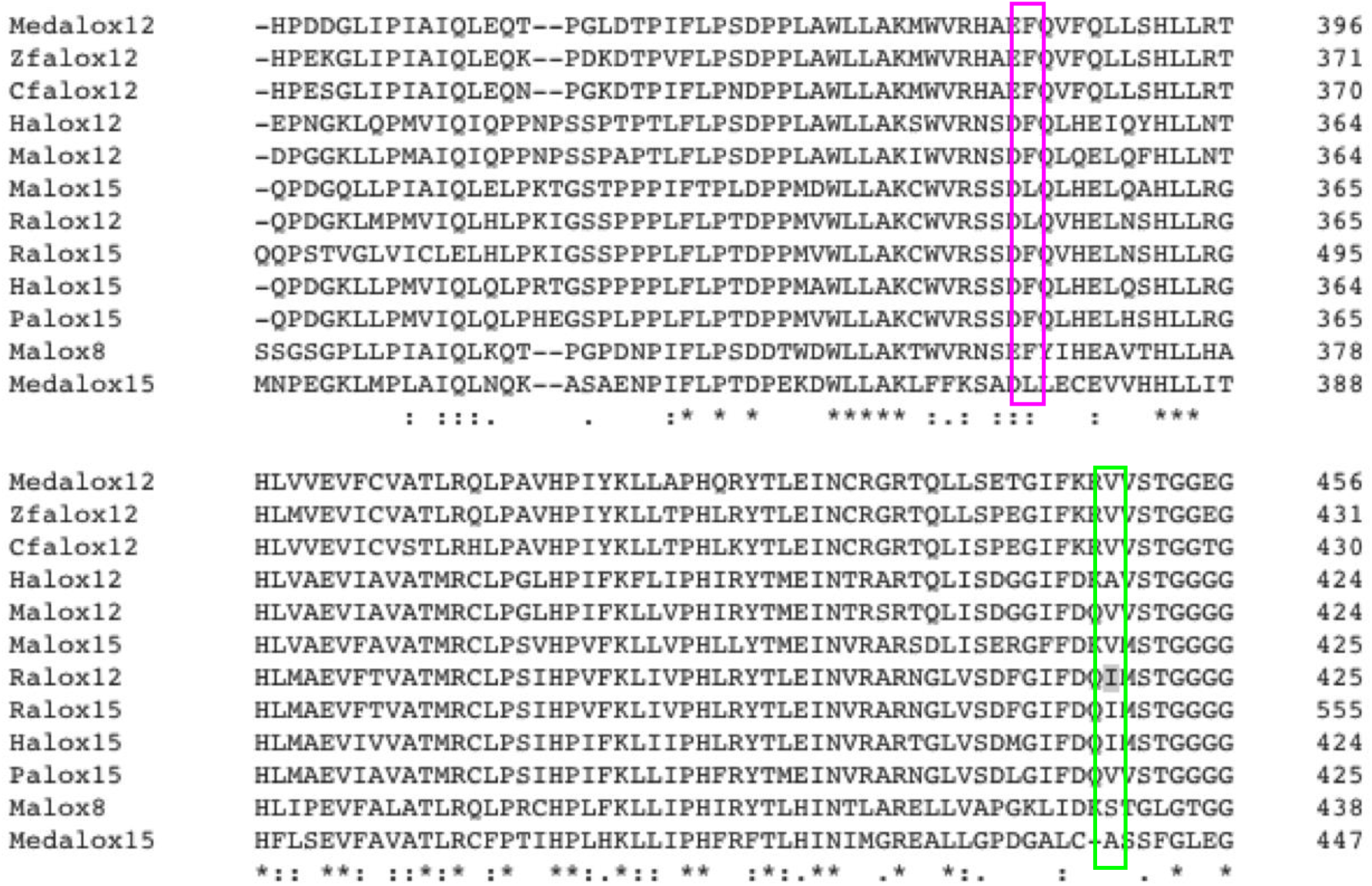
Amino acid position 353 and 418 determines LOX functionality. Alignments of mammalian and fish 12- and 15-LO proteins. A Phenylalanine **(F)** at position 353 (magenta box) in reference to rabbit 12-LO, and a bulky amino acid Valine **(V)** at position 418 (green box) dictates preferential oxygenation at carbon 15, indicating zebrafish 12-LO functions as a 15-LO lipoxygenase.

## Footnotes

1 compared to the rabbit leukocyte 12-LOX protein sequence (*67*) and associated with predominantly 15-LO activity in mammalian lipoxygenases (*68*).

